# A quadratic analytical solution of root pressure generation provides insights about bamboo and other species

**DOI:** 10.1101/2021.12.21.473687

**Authors:** Dongmei Yang, Xiaolin Wang, Mengqi Yin, Yong-Jiang Zhang, Guoquan Peng, Melvin T. Tyree

## Abstract

We derived a steady-state model of whole root pressure generation through the combined action of all parallel segments of fine roots. This may be the first complete analytical solution for root pressure, which can be applied to complex roots/shoots.

The osmotic volume of a single root is equal to that of the vessel lumen in fine roots and adjacent apoplastic spaces. Water uptake occurs via passive osmosis and active solute uptake (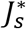, osmol s^−1^), resulting in the osmolal concentration *C*_*r*_ (mol·kg^−1^ of water) at a fixed osmotic volume. Solute loss occurs via two passive processes: radial diffusion of solute *K*_*m*_ *(C*_*r*_-*C*_*soil*_), where *K*_*m*_ is the diffusional constant and *C*_*soil*_ is the soil-solute concentration) from fine roots to soil and mass flow of solute and water into the whole plant from the end of the fine roots.

The proposed model predicts the quadratic function of root pressure 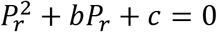, where b and c are the functions of plant hydraulic resistance, soil water potential, solute flux, and gravitational potential.

The present study investigates the theoretical dependencies of *P*_*r*_ on the factors detailed above and demonstrates the root pressure-mediated distribution of water through the hydraulic architecture of a 6.8-m-tall bamboo shoot.

**One sentence Summary:** An analytical solution for root pressure is derived and illustrated by applying it to the measured hydraulic architecture of a 7-meter bamboo shoot.

## Introduction

Typically, positive root pressure is generated in plants at night when transpiration is minimized. Positive root pressure is generated in many species, albeit not universally amongst all taxa. Generally, root pressure is the most frequent and physiologically important in monocots and woody vines (lianas), which do not exhibit secondary xylem growth in monocots or have large and vulnerable xylem vessels, respectively (Fisher et al., 1997; Ewers et al., 1997).

The putative role of root pressure is often thought to be embolism reversal on a daily or seasonal basis. Although embolism can impair xylem functionality in all species, in those exhibiting abundant secondary xylem growth, embolised vessels can be replaced seasonally through new xylem vessel growth. Root pressure may be the only mechanism to deal with embolism on an annual or even a daily basis (Cao et al., 2012) in perennial monocots, such as bamboos, with lifespan of many years.

Herein, we report the complete analytical solution of a quantitative model that can describe the origin and magnitude of root pressure in plants of any size. We hope that this solution will promote further research into root pressure using the proposed model that quantitatively explains how and why root pressure is generated. This model will enable the formulation of testable hypotheses and encourage more precisely directed experimental approaches.

Root pressure is often measured in excised roots by stopping the exudation of xylem sap from roots and measuring the maximum pressure (Steudle and Peterson, 1998; Henzler et al., 1999). The osmotic effects acting on a semi-permeable ‘membrane’ are implicated in the generation of root pressure. In living cells, this membrane is the plasmalemma membrane and the object stopping the water flow is the cell wall that stretches out elastically, thus applying a positive turgor pressure, *P*_*t*_, on the fluid to balance the osmotic pressure, *RTC*, within the cells and achieving water potential equilibrium, given as *Ψ*_*cell*_ = *P*_*t*_ − *RTC*_*cs*_, where *C*_*cs*_ is the osmolality of the cell sap. However, stable pressure also implies stable *C*_*cs*_, and the stable solute concentration in intact cells implies metabolic pathways that maintain the homeostasis of organic substances and membrane transport processes that maintain the homeostasis of inorganic ions via the action of active pumps and diffusion (leakage) through membranes. Note: All symbols used in this article are defined at their first mention in the text, and Table 1 provides a quick reference to all symbols used.

**Table 1.**
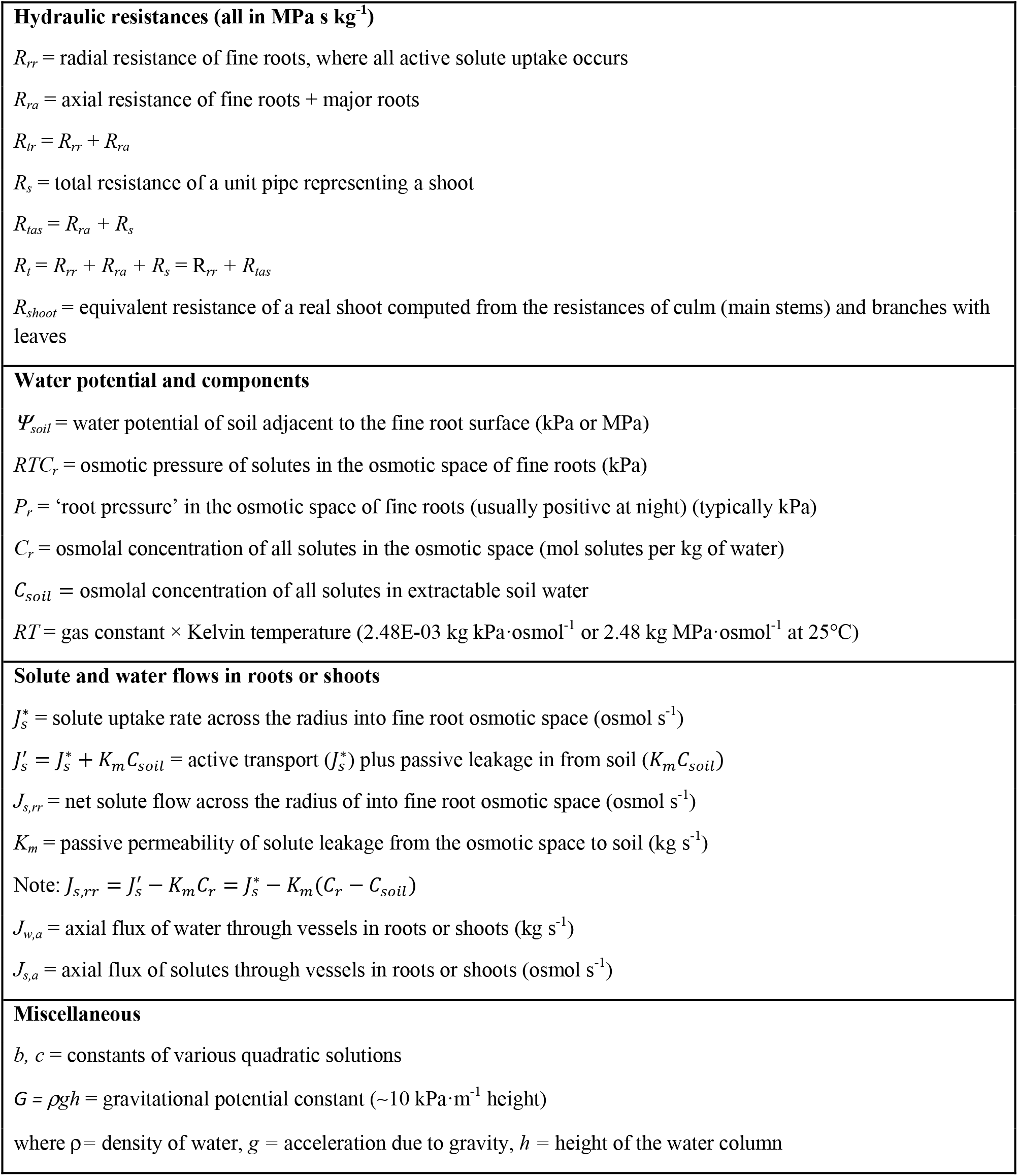
Symbols used in this article.

In fine roots, the semi-permeable ‘membrane’ is the Casparian strip, which is formed by deposits in a cell wall layer and is more impervious to solutes than water. Although previous works have speculated that neither solutes nor water can pass through the Casparian strip (e.g., Taiz et al., 2015), this is more of a conjecture rather than an experimental proof (see Geldner, 2013; Reyt et al., 2021). Thus, water flow to the stele and xylem occurs through a combination of membranes of cells in the endodermis, which includes the Casparian strip, and cell walls, although the precise pathways remain elusive. This complex tissue path permits water uptake but reduces solute leakage into the liquid phase in contact with the root surface. The low root reflection coefficient, σ_root_, reported by Henzler et al. (1999) is consistent with our tentative concept of this complex pathway.

Solute uptake decreases the solute potential, given as *Ψ*_*x,s*_ = −σ_root_ *RTC*_*r*_, where *C*_*r*_ is the osmolal solute concentration in the vessels of fine roots; *Ψ*_x,s_ further reduces the xylem water potential, thus drawing in water and generating root pressure, which is then distributed throughout the plant or its cut segment. Gravity and hydraulic pressure in the roots or shoots impact water uptake. If the root system is excised from the plant and the flow is stopped mechanically, then the maximum root pressure is achieved, much like in a single cell. Therefore, the osmotic pressure *RTC*_*r*_ reaches the maximum value during solute transport into the xylem, which increases *C*_*r*_, and a stable maximum value can be achieved when the active rate of solute uptake, *J*\*_*s*_, is balanced by the equal rate of solute loss, *J*_*s,rr*_. Notably, σ_root_, whose value is <1, is often referred to as the reflection coefficient of a fine root; in reality, however, it is the product of the reflection coefficient and osmotic coefficient (Nobel and Sanderson, 1984; Steudle and Peterson, 1998). In a system with more than two components (1 solute species and water), the meaning of the reflection coefficient is not well defined or understood in our opinion. In the series of equations described below, σ_root_ is omitted, but it can be reinstated wherever *RTC*_*r*_ appears.

## Theory and Model Equations

Figure 1 shows a schematic of the branching structure of a typical bamboo shoot. A typical ~7-m-tall bamboo plant (*Bambusa textilis*) bears 20–30 nodes with side branches, whereas a 7-m-tall bamboo shoot bears over 30 nodes, as some of them have no side branches. First, we modelled the physiology of the bamboo root system using a single root attached to a standing pipe with a root section and a pipe section above the ground (Fig. 1 far right). The root and pipe were scaled to produce the same resistance as would a large bamboo plant. All the derivations in this paper and supplemental sections are for steady state flow, which by definition does not include storage flow.

**Figure 1.**
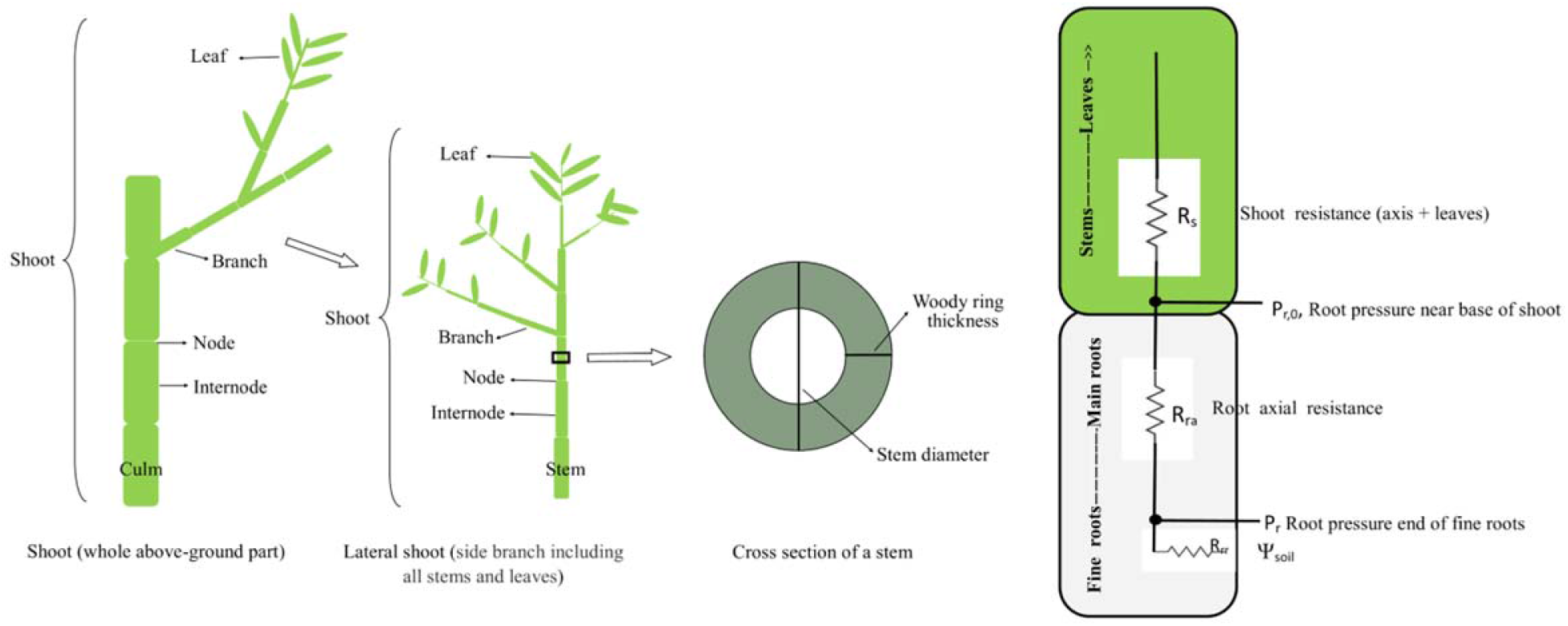
Schematic diagram of the above-ground structure of bamboo showing different components. On the far right is the resistance diagram for the root model consisting of *R*_*rr*_, fine root radial resistance; *R*_*ra*_, total axial root resistance; *R*_*s*_, shoot resistance, in a 7-m tall bamboo shoot Rs is replaced with a more complex shoot resistance diagram.

Below is the derivation of a quadratic solution for root pressure, *P*_*r*_, and root osmotic pressure, *RTC*_*r*_, where *RT* is the absolute temperature times the gas constant. *P*_*r*_ is defined as the fluid pressure at the base of the fine root, where it connects to the major root system. At this junction, all active solute uptake is stopped; thus, beyond this point, the root pressure is *P*_*r,x*_ at distance *x* from the minor root. *P*_*r,x*_ progressively decreases as *x* increases; when the solution flows further up the plant, *P*_*r,x*_ decreases to overcome gravity and hydraulic resistance. The solution of the series of equations below depends on specific ‘rate constants’ that limit the rate of water and solute flow in fine roots, where, by definition, all water and solutes are absorbed. The water flow rate is determined by two hydraulic resistances (MP_a_ s kg^−1^) in series

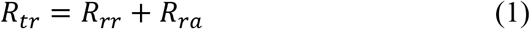

where *R*_*tr*_ = total root resistance, *R*_*rr*_ = radial root resistance (semi-permeable), and *R*_*ra*_, axial root resistance.

In terms of the equivalent resistances of the complex root and shoot pathways of water flow through the whole plant resistance, *R*_*plant*_ can be given as *R*_*tr*_ + *R*_*s*_.

Solute fluxes must be defined at two places: (1) radially in or out of the fine roots and (2) axially beyond the fine roots. The active radial flux of solutes into the root radius is 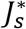 (in osmol s^−1^), and the passive radial efflux of solutes out of the fine root is *J*_*s,rr*_ (in osmol s^−1^). The efflux or leakage is determined by the passive conductance of solute along the root radius *K*_*m*_ (in kg s^−1^) and the solute concentration *C*_*r*_ (in osmol m^−3^). *J*_*s,rr*_ is calculated as the sum of solute leakage from the xylem tissues of all fine roots in the whole plant (in osmol s^−1^):

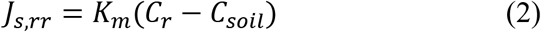

The solute flux is further driven by the water solution flow. The axial transport of solute *J*_*s,a*_ (in osmol s^−1^) can be given by Eq. (3):

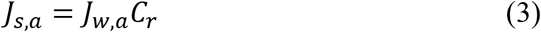

where *J*_*w,a*_ is the axial flow of water in kg s^−1^

The conservation of solutes requires the active flux into fine roots to be equal to the sum of the passive radial efflux out of the fine roots and the solute flux into fine roots and stems beyond the fine roots. The equation for the conservation of solute can be given as follows:

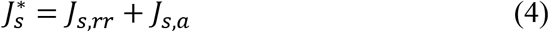

The radial flow of water into the fine roots is driven by ‘osmotic’ water uptake made possible by the semi-permeable nature of the Casparian strip. Thus, if *J*_*w,rr*_ = radial flow of water, *RT* = gas constant × Kelvin temperature = 2.48 kg MPa osmol^−1^, *P*_*r*_ = hydrostatic pressure in the fine roots (in MPa), and *RTC*_*r*_ = osmotic pressure in the root (in MPa) and if we assume *Ψ*_*soil*_ = 0, then

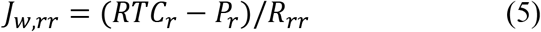

In the xylem of the root axis, water and solute flow is driven solely by pressure.

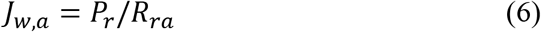

Gravity will produce any effect when root pressure is high enough to allow axial flow, which occurs when *P*_*r*_is more positive than the gravitational potential *ρgh*, where *ρ* = the density of the solution (in kg m^−3^), *g* is acceleration due to gravity (9.8 m s^−2^), and *h* is the height to the point of interest on the plant. However, we will delay the discussion on the impacts of *ρgh* and *Ψ*_*soil*_ until after the full system of equations for water flow through roots and shoots is presented. For now, we will assume that the soil water potential is zero and that there is no effect of gravity. No gravitational effects would correspond to horizontal root and shoot systems.

The value *P*_*r*_ in all equations represents pressure at the base of the fine roots; by definition, fine roots are the site where all processes of solute and water uptake occurs. Since our theory of root pressure involves exudation from whole shoots or lower parts of shoots, the pressure at the end of the vessels in the flow pathway is zero, that is, equal to atmospheric pressure. Because pressure is usually defined relative to the atmospheric pressure, there is no need to specify the ‘exit’ pressure in equations that contain *P*_*r*_. The purpose of the derivation is to obtain a quadratic solution for *P*_*r*_ in terms of the known parameters, such that *P*_*r*_ and *C*_*r*_ can be calculated given the input parameters *K*_*m*_, *R*_*ra*_, *R*_*rr*_, and 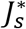; then, the application of Ohm’s law allows the calculation of *P*_*x*_ at any points in the xylem of the root and shoot.

Solution flow (mostly water) is given in kg s^−1^, and the conservation of mass requires *J*_*w,rr*_ and *J*_*w,a*_ to be equal; hence, using Eqs. (5) and (6), we can solve the equation for *RTC*_*r*_ as follows:

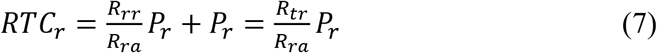

where *R*_*tr*_ = *R*_*rr*_ + *R*_*ra*_

From the conservation of solutes, Eq. (8) can be obtained for fine roots, where solute and water are absorbed; hence, we expect that the following equation to be true beyond the absorption zone in minor roots:

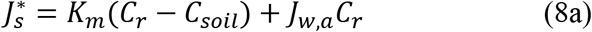

Let

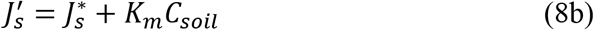

Then

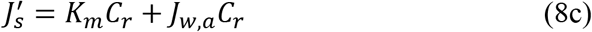

where *K*_*m*_(*C*_*r*_ − *C*_*soil*_)is the passive loss of solutes from the root radius to the soil and *J*_*w,a*_*C*_*r*_ is the axial transport of solute as water flow pushes them out beyond the fine roots. The passive fluxes could be viewed as two independent influx and efflux components, *K*_*m*_*C*_*soil*_ and *K*_*m*_*C*_*r*_, respectively, where the sum of the two is the net passive flux.

In Eq. (8a), 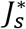 is the active uptake of solutes from the soil; there is no need to specify a distinct mechanism because the value of 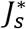 is independent of mechanism. Based on Eq. (6), we can solve (8) for *C*_*r*_ as follows:

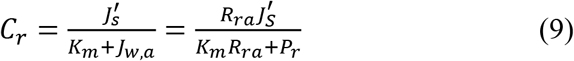

Furthermore, we can solve Eq. (7) for *P*_*r*_ as follows:

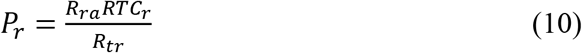

By substituting Eq. (9) into Eq. (10), the following quadratic equation for *P*_*r*_ can be obtained

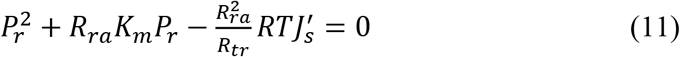

Now, the coefficient *b* and *c* can be given as follows:

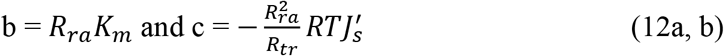

The following quadratic solution can be obtained:

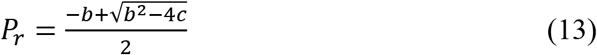

The solution for quadratic equations always has a 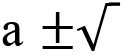 term; however, we can reject the negative alternative because it yields a negative *P*_*r*_, and the solution of interest is for positive values of *P*_*r*_, which by definition is root pressure. We have also omitted the root reflection coefficient (σ_root_) from all equations above and below, because it is not accurately known in most cases; however, readers can add it as an implied coefficient wherever *RT* occurs. If we substitute the *P*_*r*_ value from Eq. (13) into Eq.(7), the value for *RTC*_*r*_ as well as other values of interest, such as *C*_*r*_ and *J*_*w,a*_, amongst others, can be calculated.

The root pressure model presented above can be applied to other systems with the addition of the resistance of shoot, *R*_*s*_, or real shoots with complex resistance networks. It can also be generalized to include soil water potential, *Ψ*_*soil*_, and gravitational potential, *ρgh*. The value of *ρg* is approximately 0.01 MPa·m^−1^. All these derivations are provided in Supporting Information (Note S1). The solution also uses combined-values, in addition to *R*_*rr*_, *R*_*ra*_ and *R*_*s*_. The first is the ‘equivalent’ axial resistance *R*_*tas*_ = *R*_*ra*_ + *R*_*s*_, and the second is the total equivalent resistance of the whole system, *R*_*t*_ = *R*_*rr*_ + *R*_*ra*_+*R*_*s*_.

Despite using these additional resistance values, the solution remains a quadratic equation, but the coefficients *b* and *c* change. If we allow negative values of *Ψ*_*soil*_, the values of *b* and *c* can be given as follows:

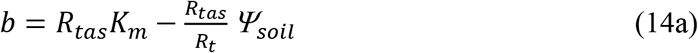

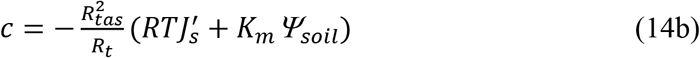

By further allowing the gravitational potential but *Ψ*_*soil*_=0, we obtain the following equation:

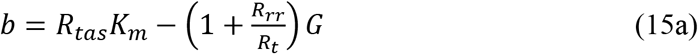

where *G=ρgh*

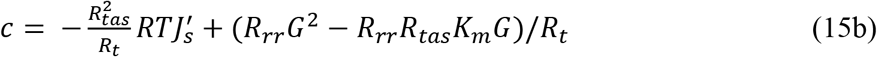

On comparing Eqs. (14) and (15) with Eq. (12), it can be observed that *b* and *c* have a similar first term with *K*_*m*_ or 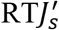 plus other terms, including *G= ρgh* or *Ψ*_*soil*_. Therefore, if G = 0 or *Ψ*_*soil*_ = 0, both model outputs return the simpler solution derived above.

An important aspect that must be considered is that all the above equations apply strictly to a very ‘linear’ plant with unbranched root and shoot systems. If *P*_*r*_ is not high enough to push water to the end of the standing pipe, that is the extension of the root, then *J*_*w,a*_ would reach zero before flowing out of the pipe, assuming the absence of nocturnal transpiration. In the context of a plant with a complex branching with leaves at different heights, water could flow out of the leaves at lower *h* values before flowing out from the higher leaves. In a plant that is taller than the height until which the root pressure can drive water, then somewhere in the branching structure, the xylem pressure will drop to 0 MPa, and beyond that point, the pressure will become negative. The cohesion tension theory explains the negative pressure caused by the radius of the curvature of water in the cell wall pores at the evaporative surface (Tyree and Zimmerman 2013). However, when the pressure exceeds zero in the lower leaves, water will flow out into the air space and/or out of the hydathodes, if present.

One may wonder whether the proposed quadratic solution for root pressure and root exudation to the top of the standing pipe can reflect a more complex situation in a real plant with leaves distributed at different heights above the ground. In the sections below, we demonstrate that the predictions of our simple quadratic solution are indeed suitable for more complex shoot structures.

In shoots, average resistance can be calculated by applying the Ohm’s law to obtain the equivalent resistance of complex shoots from the measured shoot components. Alternatively, the equivalent resistances can be measured directly in roots and shoots using an HPFM.

To scale the model to represent different plant sizes, the objective was to adjust the resistances in Eqs. (12, 14, and 15) to be equal to the resistance of smaller or larger plants and to adjust *h* to scale for taller or shorter plants. In general, because larger plants have more branches (see Yang et al., 2021), plant size is inversely proportional to resistance, that is, lower resistance in larger plants and higher resistance in smaller plants. The *K*_*m*_ value for passive radial leakage from fine roots is proportional to the root size measured in terms of fine root surface area, that is, larger values of *K*_*m*_ for bigger fine root systems, all working in parallel. Detailed information on scaling factors for *B textilis* has been reported by Yang et al. (2021).

## Model-Predicted Results

### Effects of axial versus radial root resistance

Vascular resistance from fine roots to fine veins in leaves accounts for only approximately half of the total resistance of plants to water transport (see numerous references Chapter 6 in Tyree and Zimmermann, 2013). The remainder half of the whole plant resistance is produced in non-vascular tissues of roots and leaves, where water travels for typically <0.2 mm.

Far less data are available for roots, although two excellent studies by Steudle and Peterson (1998) as well as Nobel and Sanderson (1984) have shown that the axial root resistance, *R*_*ra*_, of the entire root system is approximately equal to the radial resistance in the minor roots, *R*_*rr*_. Figure 2 shows the impacts of unequal values on the transport parameters computed by the proposed model.

**Figure 2.**
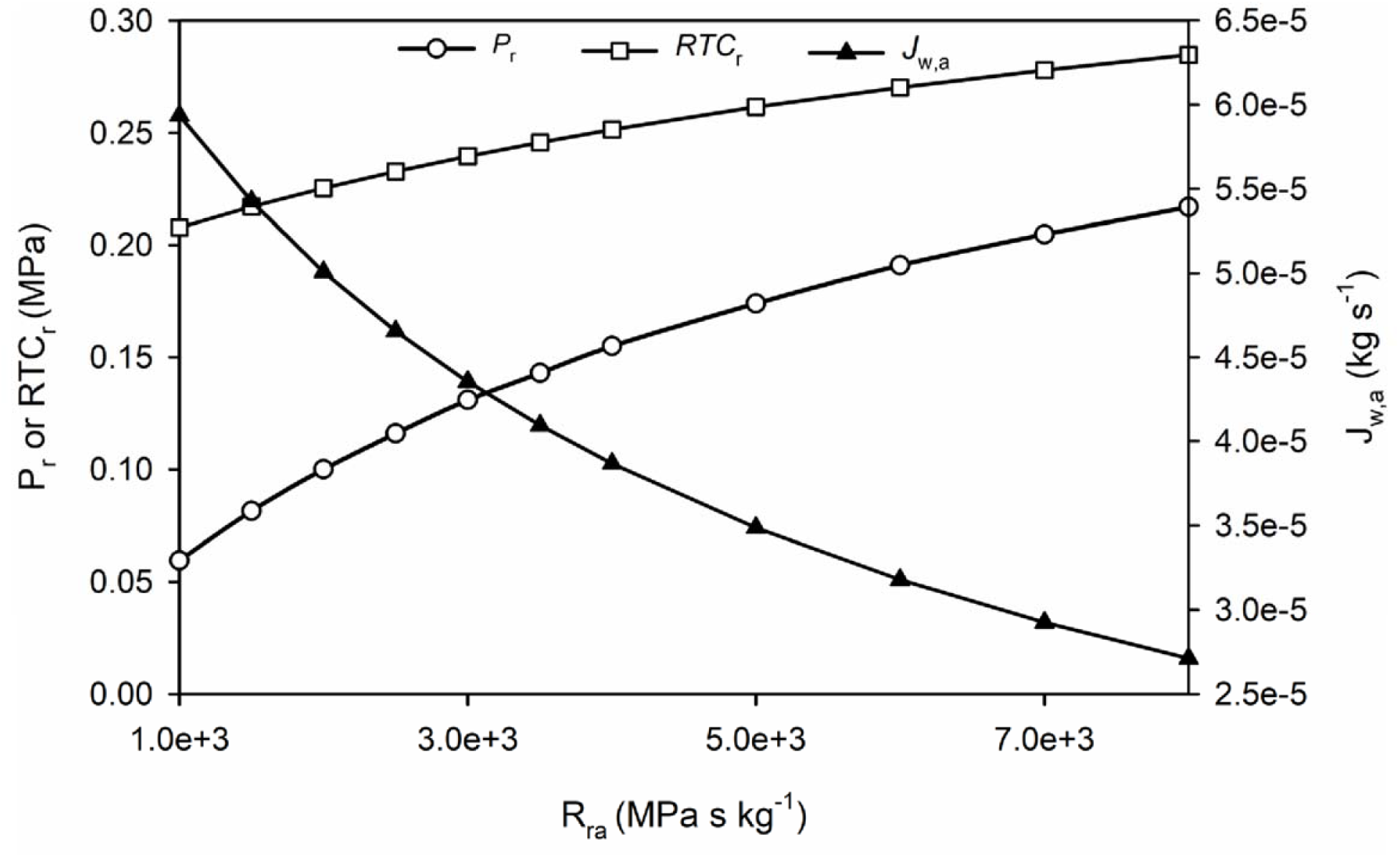
Model of an extensive root system with the radial resistance of many fine roots working in parallel, *R*_*rr*_, held constant at 2.5×10^3^ MPa s kg^−1^ and *R*_ra_ changed as shown on the *x*-axis to demonstrate the impact of the combined equivalent resistance of axial flow beyond the fine roots to the soil surface. The other parameter used were *K*_*m*_ = 6×10^−5^ Kg s^−1^ and 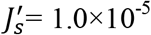 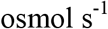.

The increase in *R*_*ra*_ reduced the sap transport rate up the axis of the main root system, *J*_*w,ra*_, which is nearly equal to the water flow into the minor roots, *J*_*w,rr*_, but increased both root pressure, *P*_*r*_, and root osmotic pressure, *RTC*_*r*_, ultimately increasing *C*_*r*_. Quantitatively, the solute concentration *C*_*r*_ depends on the ratio of net solute uptake and the water flux into the fine roots, *J*_*net,s*_*/J*_*w*,rr_. Since 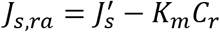, the value of 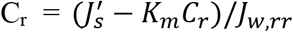. As *K*_*m*_ and 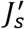 are constants, *C*_*r*_ increased with decrease in *J*_*w,ra*_. Of course, *C*_*r*_ cannot increase without limit; there is some passive leakage out of the root, given as *K*_*m*_*C*_*r*_. In our model, the value of the net solute influx was close to its theoretical limit. In Figure 2, the values of *K*_*m*_ and 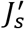 were selected such that the rise in concentration was within one-third of the theoretical maximum concentration. The theoretical maximum value of *C*_*r*_ is achieved when the flow of the solution out of the minor roots is stopped by infinite hydraulic resistance.

Further, the root pressure model was modified to examine the effects of soil water potential and *Ψ*_*soil*_ on root exudation. The *b* and *c* components are expressed in Eqs. (14a and b). The first terms for *b* and *c* are identical to those in the simple solution shown in Eq. (12); however, the *b* and *c* ‘constants’ include terms with *Ψ*_soil_. Notably, as plants grow in size, the hydraulic resistances and *K*_*m*_ constants in this simple model must be modified.

Figure 3 presents a sample plot showing the theoretical effects of soil water potential on *P*_*r*_ in the fine roots and *P*_*r,0*_ near the soil surface. Even though the values of root pressure (*P*_*r*_ and *P*_*r,0*_) fell, they decreased with a slope of <1; as such, *P*_*r*_ decreased from the initial value of 0.209 MPa to zero with a decrease of 0.400 in *Ψ*_*soil*_, yielding an average slope of 0.512. Therefore, the root pressure mechanism of the proposed model dynamically responds to soil water potential by increasing the value of osmotic pressure, as the passive water flux, *J*_*w,rr*_, into fine roots falls faster than the net solute influx, *J*_*s,rr*_ (data not shown). Hence, the root pressure remained higher than it would have been if the roots had not dynamically responded to changes in soil water potential. This adjustment in the water relations of roots is evident from the rise in osmotic pressure with the fall of soil water potential.

**Figure 3.**
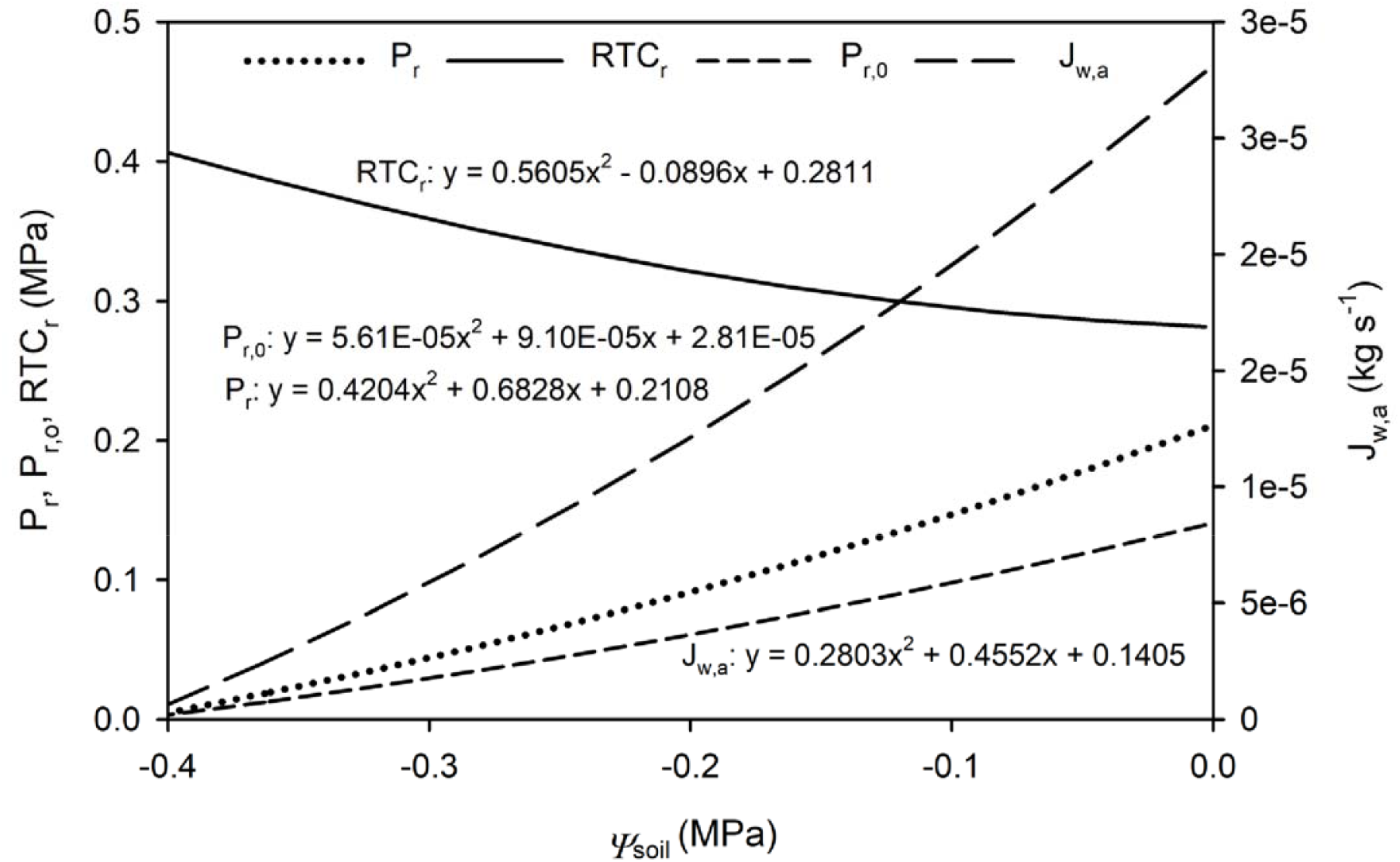
Theoretical impact of soil water potential (*Ψ*_*soil*_) on root osmotic pressure (*RTC*_s_), root pressure at the end of the fine roots (*P*_*r*_), root pressure at the soil surface (*P*_*r,0*_), and root pressure-driven sap exudation (*J*_*w,a*_). *R*_*rr*_ = *R*_*ra*_ = 2.5×10^−3^; *R*_*s*_ = 5×10^−3^; *K*_*m*_ and 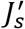 as in Figure 2.

The next logical aspect to examine was the impact of gravitational potential on the dynamics of root pressure generation, as predicted by our simple model. Again, we used a simple unit pipe structure for roots and shoots but scaled these with hydraulic resistances that approximate bamboo roots and shoots. Therefore, the equivalent resistance of the shoot was measured with a >11-m-long flexible pipe, the end of which could be lifted to any height, *h*, from the ground level (*h* = 0 m) to a maximum of 10 m above the ground. This rise in height corresponds to the change in gravitational force from 0.0 to 0.1 MPa. The derivation is shown in the Supporting Information (Note S1), and the addition of gravitational potential entailed the incorporation of several other terms to the ‘constants’ *b* and *c* [see Eqs. (15a and b)].

The proposed root pressure model appeared to respond dynamically to changes in the gravitational potential *G*. With rise in *G* to 0.1 MPa, the osmotic pressure increased to compensate for the 26% increase in *G.* The root pressure values *P*_*r*_ in the fine roots and *P*_*r,0*_ at the ground level increased at slower rates but still reduced the impact of gravitational potential by 44% and 30%, respectively (Figure 4).

**Figure 4.**
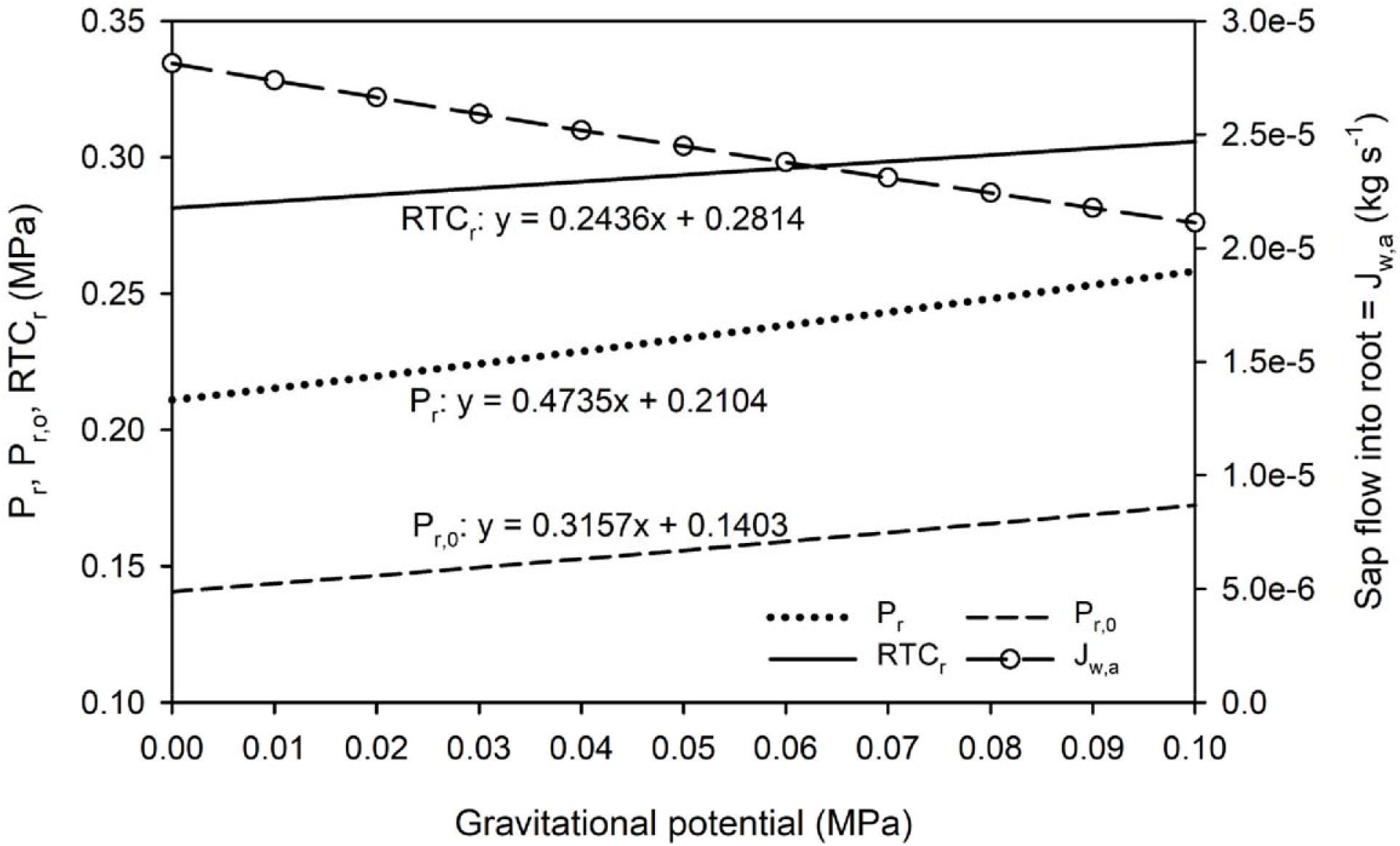
Theoretical model predictions for the generation of root pressure and response to changes in gravitational potential (*ρgh*). The model parameters used were scaled to those of a tall bamboo shoot. 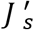 (mole s^−1^) is the solute uptake rate; *K*_*m*_ (Kg s^−1^) is the passive solute permeability constant; and *R*_*rr*_, *R*_*ra*_, and *R*_*shoot*_ are the hydraulic resistances of fine roots, whole root axis, and whole shoot axis, respectively (all in MPa s kg^−1^). 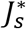, *K*_*m*_, *R*_*rr*_, *R*_*ra*_, and *R*_*s*_ as in Figure 3. For this figure, h = 0 (the junction between fine and major roots).

Consistently, this trend was attributed to the reduction in the flow of water into and out of the fine roots (solid line with points in Figure 4), which increased the ratio of net solute flux/water influx to *C*_*r*_ and ultimately the osmotic pressure *RTC*_*r*_. Of note, we do not claim that the proposed simple model reflects real scenarios. Nonetheless, this model is the first quantitative solution that predicts the dynamic nature of root pressure and could thus be employed in future experimental trials.

In the bamboo shoot, root exudate can be observed coming out of many leaves at different heights. Therefore, we evaluated the predictive performance of the proposed model using a hydraulic system that is comparable to a tall bamboo shoot.

### Root pressure in a tall bamboo shoot

Recently, Yang et al. (2021) reported detailed hydraulic architecture of bamboo plants (*B. textilis*) varying in height from approximately 2 to 7 m. The authors reported the relationships of the resistance of the main stem (culms), side branches, and leaves scale with plant size. From their study, we obtained the dataset for a 6.8-m-tall shoot to compute the equivalent resistance of each of the 25 nodes bearing leaves from the base to the apex of the main culm using the Ohm’s law. Using these data, we calculated root pressure at each shoot-bearing node and the flow rate out of the node to all side branches that bear a known leaf area based on measured branch resistances. Once the magnitude of the root pressure at the base of a shoot was known, the water flow rate through all the nodes towards the outside as well as that through the leaves could be calculated, as long as the component resistances in the branched flow path were measured. For more details, see Supporting Information (Note S2, Model S1).

During the monsoon season, water can be seen dripping from all leaf margins (when it is not raining) of *B. textilis*. Here, root pressure values were not measured. Instead, using the hydraulic model above, we calculated the possible root pressure that could induce exudation from these tall bamboos and confirmed that the root pressure was within the range of pressures measured by Cao et al. (2012) and others for 24 bamboo species of various heights.

The model reported in the present study is based on the measurements for an individual 6.8-m-tall bamboo with known leaf distribution, as shown in Figure 5. The total leaf area serviced by the culm at different heights followed a smooth sigmoid curve, indicating that there was less leaf area at the base and the apex of the bamboo shoots, as evidenced by the values of leaf area per node in the inset of Figure 5. Actual measured values of leaf area were used in the model.

**Figure 5.**
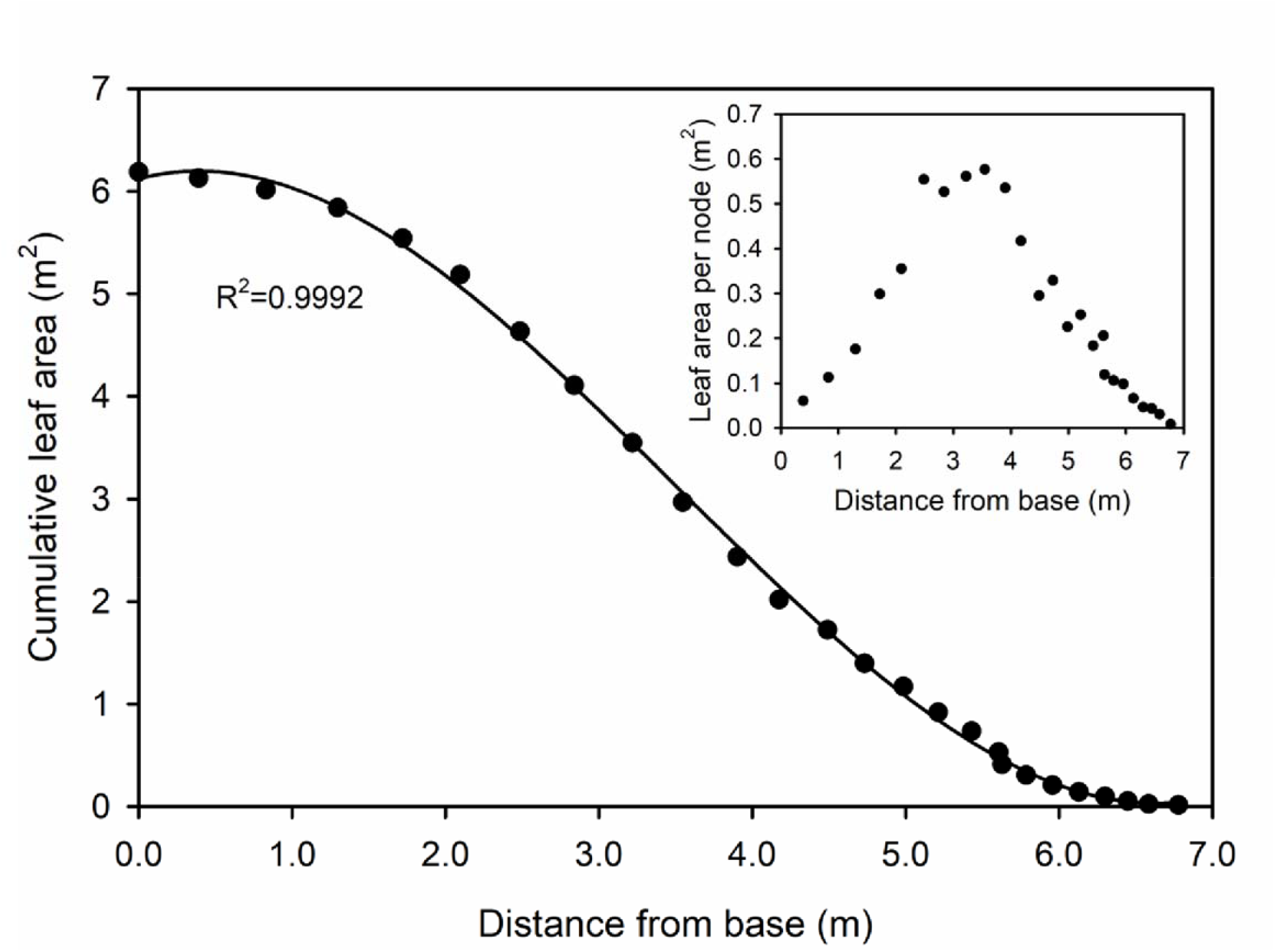
Leaf area serviced by the main stem (culm) *versus* distance (height) up the stem. The inset shows the leaf area attached to the side branches at each node *versus* distance.

Figure 6 presents a plot of the resistance of the culm and side branches + leaves *versus* height up 6.8 m. The typical resistance of shoots (for water flow from the nodes to the leaves) was fairly independent of nodal position (see Figure 5 in Yang et al., 2021) when scaled to the leaf area of 4×10^4^ MPa m^2^ s kg^−1^; therefore, the absolute resistance was inversely proportional to the leaf area *A*_*L*_ (i.e. 4×10^4^*/A*_*L*_ in MPa s kg^−1^). At the base of the bamboo shoot, water passes through many nodes before encountering side branches with nodes; however, within approximately 2 m of the apex, every node bears at least few side branches. Conversely, in the vertical bamboo culm, there was very small hydraulic resistance along most of the length of the shoot until the apex. Overall, the hydraulic resistance distribution of bamboo shoots was similar to that of all other tree shoots studied thus far. The main resistance to water flow occurred in minor twigs and leaves. In bamboo, the hydraulic resistance of the leaf-bearing branches and leaves was two orders of magnitude higher than that of the culm segment, with most of the resistance being confined to the leaves.

**Figure 6.**
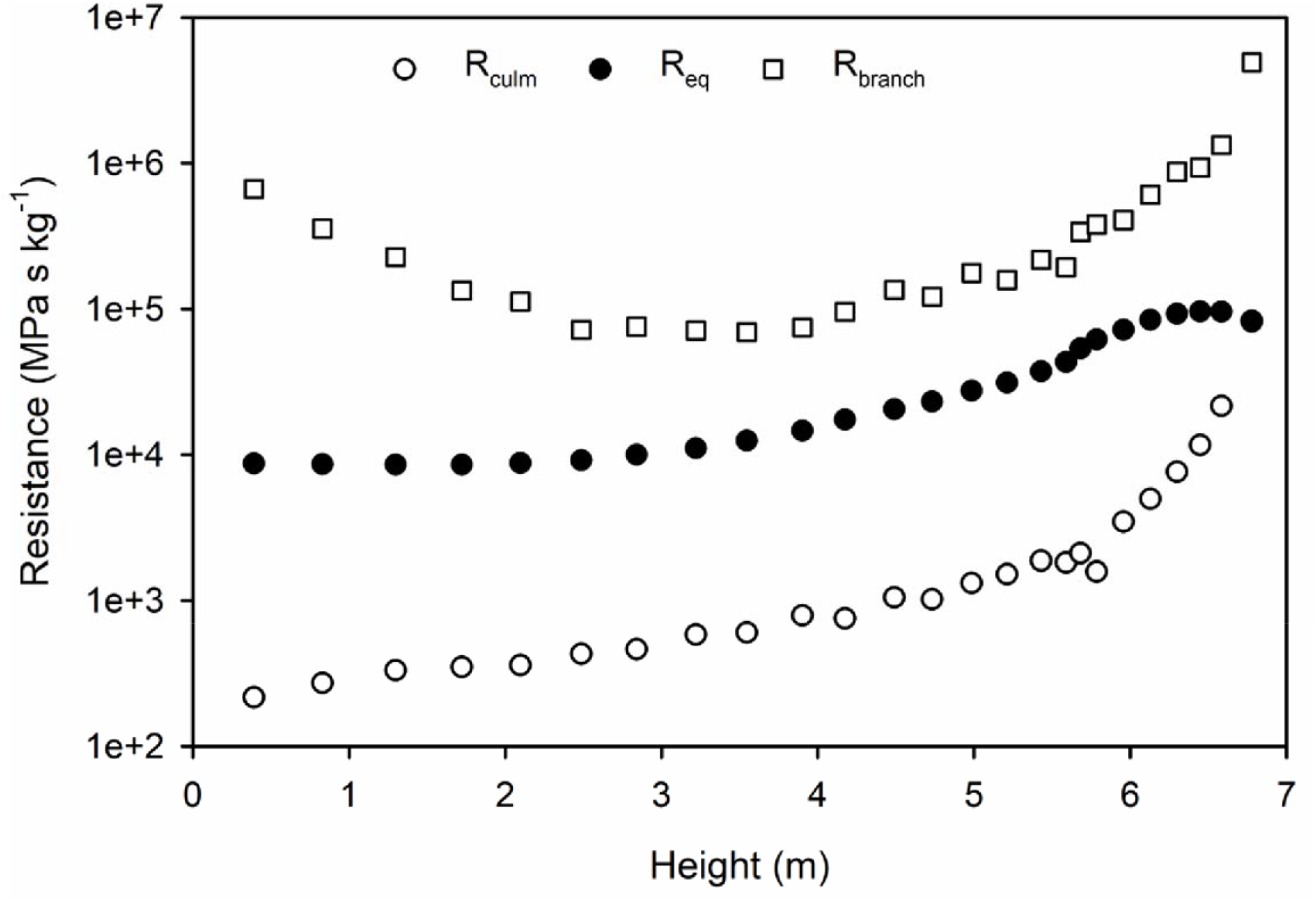
Changes in hydraulic resistance with distance from the base to the apex of shoot. R_culm_, hydraulic resistances of culm segments between nodes with side branches. Not all nodes bear side branches. R_branch_, hydraulic resistances of side branches + leaves. R_eq_, the equivalent resistance of the entire shoot from the base to leaves at each height.

The computed pressure gradient up the vertical culm is shown in Figure 7. The pressure at the base of the shoot exceeded the gravitational potential line (*G* = *ρgh*) by approximately 0.094 MPa. The pressure difference required to propel the xylem sap upwards and out of the leaves is called the hydrokinetic pressure difference (*P*_*hk*_), and it is indicated by the height of the double arrowheads in Figure 7. A large value of *P*_*hk*_ is required to push water through the stem and leaves, starting at the node and ending at the leaf margins (=atmospheric pressure), where hydathodes are present, or in the internal leaf air spaces. The value declines towards the apex of the culm whilst still remaining above the *G* line.

**Figure 7.**
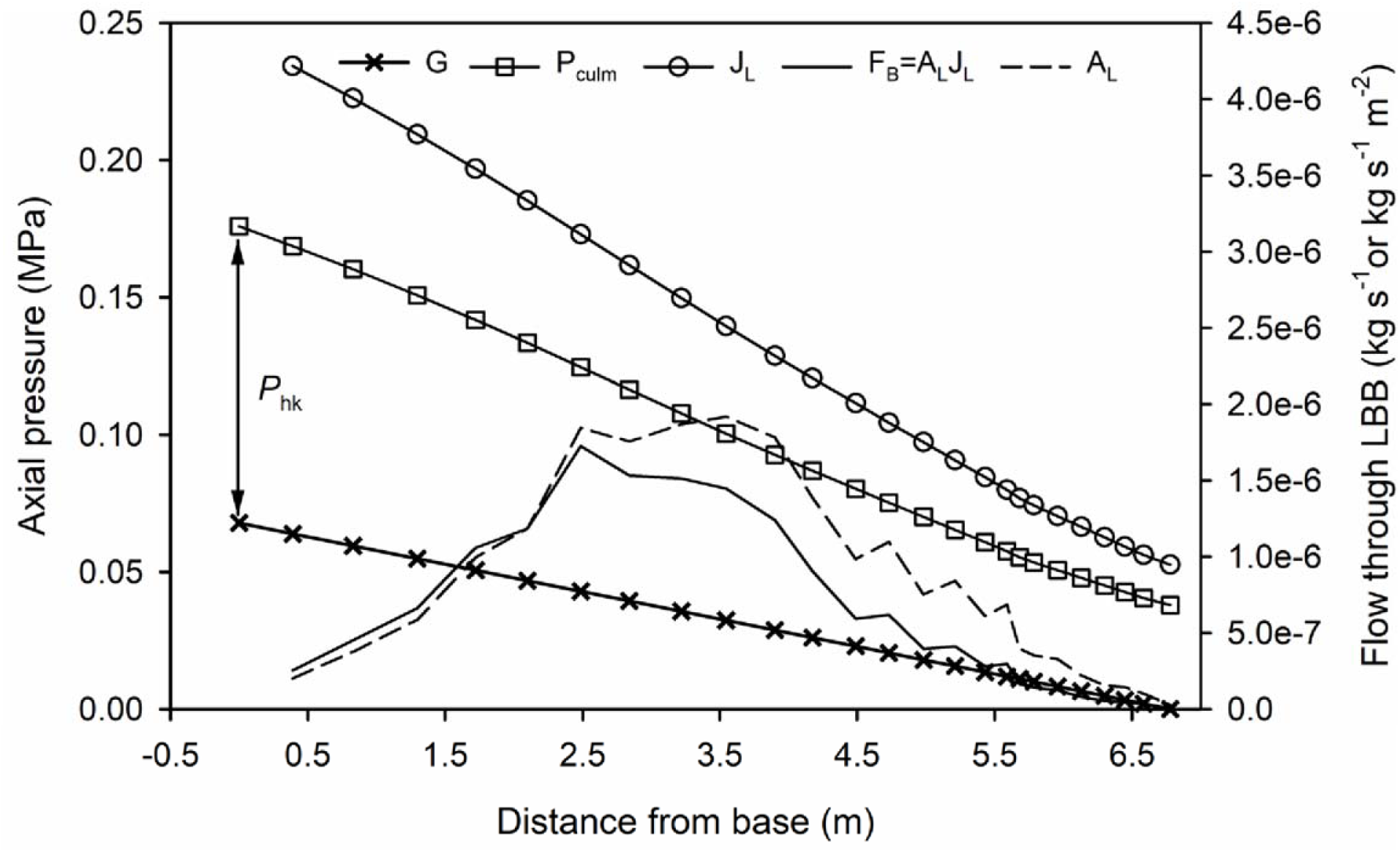
Pressure gradient up the vertical culm of bamboo. On left ordinate axis, pressure profile in culm (*P*_*culm*_) and gravitational potential (*G = ρgh*) are shown. On the right ordinate axis, flow through leaves per unit leaf area (*J*_*L*_ *kg s*^−1^ *m*^−2^) and absolute branch flow (*F*_*B*_, kg s^−1^) *versus* distance from the base for a vertical culm on the *x*-axis are shown. The area of leaves (*A*_L_) was scaled (divided by 3×10^5^) such that it could be displayed on the left ordinate axis. Note: *P*_*hk*_ is the hydrokinetic pressure difference (=double arrows) and is equal to the difference between *P*_*culm*_ and *G* at each height. Values of resistances were scaled to match the R_shoot_ of bamboo. LBB, leaf bearing branch. *R*_*rr*_ = *R*_*ra*_ = 4.337×10^−3^, *R*_*shoot*_ = 8.673×10^−3^, *K*_*m*_ = 3×10^−5^; 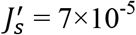.

Our simplified root model describes the shoot resistance as a unit pipe with all the sap flowing out the end at 6.8 m height, whereas sin Fig. 7 the contribution of *G* is computed at the height of each node. In an intact root connected to the shoot, the processes are more intricate. If water is pushed up against the gravitational potential gradient, then the restoring pressure *ρgh* is generated, where *ρ* is the density of xylem sap, *g* is acceleration due to gravity, and *h* is the height above or below a reference point at which the pressure is measured. The value of *ρg* is approximately 0.01 MPa m^−1^. In the presence of root pressure, water is often seen flowing out from the leaf margins, where hydathodes are present, from stomatal pores on the underside of the leaf, or from the xylem of any recently injured leaf or stem. Therefore, the observed root pressure is expected to be equal to the combined impact of gravity and hydrokinetic resistance to flow and can be given as *R*_*h*_*J*_*w*_, where *J*_*w*_ is the flow of the solution in any root or stem system and *R*_*h*_ is the hydraulic resistance of the flow of the whole system (root+shoot). This is analogous to the Ohm’s law, V = IR, where voltage (V) is analogous to root pressure, current (I) is analogous to water flow, and electrical resistance (R) is analogous to hydraulic resistance; however, a major exception to this analogy is that electrical circuits are not influenced by gravity. The flow pathway and resistance in the root and shoot systems are formed by a complex network of stem or root segments, each with a finite resistance, *R*_*i*_; however, any complex resistance network can be reduced to an equivalent resistance network, given as *R*_*eqv,rt*_ or *R*_*eqv,sh*_, where the subscript *rt* refers to the whole root and *sh* to the whole shoot. The values of equivalent resistance can be quantified using a high-pressure flow meter (HPFM) or calculated theoretically based on the measurements of serial and parallel hydraulic resistance elements through repeated use of Ohm’s law.

About 2.5 m up the culm, the rate of solution flow into the side branches reached the maximum; however, the maximum rate in the leaf area attached to the side branches was reached at 3.55 m. The solid black line in Figure 7 indicates the net flow of the solution (kg s^−1^) into each leaf-bearing branch. The root pressure required to produce these profiles when measured at ground the level (*x* = 0) was close to the range reported for some other bamboo species (up to 0.16 MPa) (Cao et al., 2012). A qualitative confirmation that the root pressure generated in *B. textilis* is indeed large enough to push water out of the leaves was obtained based on the observation of copious extrusion of water from the hydathodes at leaf margins, even high up in the shoot. Notwithstanding, there would be a large amount of pressure to spare. Even at a root pressure of ~0.096 MPa, the stem pressure would remain above the value of *G* throughout the length of the culm >6 m tall and would drive water into all side branches and leaves (data not shown).

The distance from the base to the apex of the bamboo culm shown in Figure 7 is also the height *h*. The model assumed that the culm was straight and vertical. However, culms are usually slightly curved or, in some cases, form a graceful arch with a maximum height of approximately 6 m up to where the culm is horizontal and then gently curves downwards; therefore, the gravitational effect may be weaker than that shown in Figure 7, and more water would reach the leaves of a curved culm.

Initial solutions violated the conservation of mass. In other words, the mass of water extruded from the top of the unit pipe model differed from the sum of water extrusion calculated from all leaves in the resistance network (*ΣJ*_*L*_) used to simulate the bamboo shoot; specifically, we noted an error of <4% of the unit pipe value, and the results varied depending on *K*_*m*_ and 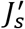. As shown in Figure 4, variations in *G* at a constant shoot resistance explain the observed violation of mass conservation.

The average *G* values experienced by leaves at each height in the bamboo shoot were different from those in the standing pipe, where all water flowed out from only one place, where the *G* value was applied. Therefore, we used the Excel solver function to study the impact of *G* on the solutions of the standing pipe *versus* those of the actual bamboo shoot. By adjusting the height of the standing pipe to be 0.17 m shorter than the 6.78-m-tall bamboo shoot conserved the mass without changing the resistance.

## Discussion

The theory described above is very simplified. For instance, no particular solutes in the xylem sap of the model roots are specified. In reality, there are a multitude of solutes with different *K*_*m*_ values and different active fluxes 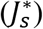. To make the proposed model more ‘realistic’, we must obtain many unknown rate constants for deriving an analytical solution. To gauge the complexity of the real-world scenarios, we refer readers to William Pickard’s work. In a previous study, Pickard (2003a) has provided a full description of theoretical events in root–soil systems, divided into three compartments of fixed volume, namely soil, symplasm, and vessels/wood. One of the purposes of Pickard’s theory was to determine how rapidly the three-compartment system responds to step changes in conditions (concentration and pressure). In another experimental study on excised tomato roots, Pickard (2003b) measured the response time to quick changes in soil osmotic potential induced by alternating irrigation with water or concentrated salt solutions. The response of the system was rapid enough (<100 s) to justify our simpler two-compartment model. Our model can be used to quantify events ranging from a timeline of 200 s to days.

Presumably, our simplified solution provides the first complete answer to how root pressure and solution flow propagate up the root axis and what happens as water is dispersed into a branched shoot system, such as a tall bamboo shoot (7 m). Moreover, our model showed how hydraulic resistance in roots and shoots affects changes in pressure and solution flow rate in a complex shoot structure. Our simplified model advances theoretical knowledge and also offers foundational data to devise experiments for testing the reliability of the theory. Such experimental trials would pave the way for creating more reliable and accurate models in the future, which focus on the complete set of events in the whole plant. In addition, our proposed model can be used to study the optimization of hydraulic architecture to maintain a relatively stable leaf water supply and selecting force to shape the hydraulic system in monocots (Yang et al., 2021).

Notably, our theory of root pressure propagation in the whole plant is parallel to the theory of phloem translocation based on the Münch pressure flow hypothesis. Tyree et al. (1974) provided the first steady-state solution of this problem, and Christy and Ferrier (1973) offered the first dynamic-state solution. Although the previous studies had boundary conditions, Goeschl et al. (1976) explored alternative boundary conditions in subsequent studies, and advances are still underway (e.g., Thompson and Holbrook, 2007; Sellier and Mammeri, 2019). The major difference, however, is that in the Münch pressure flow model, the water and solutes pass radially through the plasmalemma membranes of sieve tubes (and/or companion cells), and in many cases, active pumps for sugars and ions are involved in both solute loading and unloading (Rennie and Turgeon, 2009; Turgeon, 2010). On the contrary, in our root pressure model, active loading of solutes occurs only into xylem of minor roots and all flow afterwards is passive. As sap flows upwards passively, growing tissues can upload mineral nutrients needed for cell enlargement, but the model does not need to account for this.

We specifically refer to the Münch pressure flow hypothesis in this study, because sap flow in the living phloem cells contains both carbohydrates from photosynthesis and minerals and nutrients from living sieve tubes and companion cells (Hayashi and Chino, 1986). These solutes must be unloaded near the growing root tips together with the carbohydrates, and then the minerals and nutrients travel back up the xylem stream to be reloaded in sieve tubes in photosynthetically active leaves. The proportions of nutrients coming from sieve tubes and soil to travel up the xylem remain unknown and warrant further research. We hypothesize that half or more of these nutrients are derived from the phloem tissue.

Furthermore, there are some important quantitative differences between xylem transport at night via root pressure and daytime transpiration. According to the cohesion tension theory, the pressure is predominately negative in the xylem fluid during daylight hours, and this negative pressure is generated at the evaporative surface. At the cell wall, the radius of the curvature of pores near the internal leaf air spaces is adjusted to produce a fluid pressure that is more negative than the atmospheric pressure. The pressure generated in the fluid is −*τ/r*, where *τ* is the surface tension of water (0.072 N m^−1^) and *r* is the radius of the curvature of the air–water interface. The value of *r* has a wide range (10^−4^ to 10^−8^ m), producing a negative pressure anywhere from 0 to −7 MPa. However, at pressures below −1 or −2 MPa, stomates start to close depending on the species, which limits the magnitude of negative pressure except when soils become very dry. Generally, stomatal closure occurs right before the xylem pressure becomes negative enough to induce a >5% loss of hydraulic conductivity in the xylem due to vessel embolism (Martin-StPaul et al., 2017; Hochberg et al., 2017). Water flow at night is driven by the pressure difference between roots and leaves, although the magnitude of flows is 10 to 30 times lower during the night than during the day (Cirelli et al., 2015). Regarding the root pressure-driven flow, as shown in Figure 7, the flow per unit leaf area was the highest near the base and the lowest at the top of the bamboo shoots. The primary difference is that root pressure-driven flow at night follows the pathway of the least resistance to leaves, whereas tension flow follows the pathway of the highest demand, that is, leaves with the highest irradiance.

In the proposed simplified model, we ignored the water evaporation rate from leaves at night, because during the rainy season in subtropical climates, where our field study was conducted, the night time humidity is often 100%. Therefore, evaporation is zero on cloudy nights, and even on nights with clear skies, the leaf temperature is below the ambient temperature, leading to dew formation on leaf surfaces. However, even under little transpiration, the benefits of root pressure can be evaluated at a leaf vessel pressure of 0 MPa. Assumedly, embolism commonly occurs in the vessels of leaves and minor branches supporting the leaves. Would zero fluid pressure be sufficient to reverse embolism? The detailed answer to this question is beyond the scope of the present model; however, based on our findings, atmospheric pressure is indeed sufficient to reverse leaf xylem embolism within <2 h, assuming that membranes do not virtually impede gas diffusion (Subczynski et al., 1992). We currently have a quadratic solution for root pressure when Ψ_leaf_<0 but this will be presented together with an experimental paper addressing root pressure with night-time transpiration.

Although the model is simplified, it can be used to design future experiments that could help refine the model. For example, in the context of a 7 m tall bamboo shoot, the shoot could be excised at night when root exudation is exhibited. The equivalent hydraulic resistance of the shoot could be measured with a HPFM. Then the HPFM connector could be attached to the root stock and thus connected to different capillary tubes with a range of resistances bigger or smaller than the resistance of the shoot. The connection device could be designed to also hold a pressure transducer. This experiment would make it easy to see how the root pressure changes with shoot resistance as shown only theoretically in Fig. 2. Alternatively, the root stock would be reconned with a 11-m-long water-filled flexible pipe and a pressure transducer at the bottom; then the end of the pipe could be raised or lowered over a 10-m range of heights to simulate changes in *G* and to monitor changes in root pressure as is shown only theoretically in Fig. 4. If the data from any of the experiments do not agree with the model, then the next step would be to modify the model to make it more predictive of the observations.

## Conclusions

The present study proposes the first model of root pressure with a complete analytical solution. The proposed model can be used to calculate the root pressure driven-flows based on the known values of hydraulic resistance, solute uptake rate 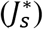, and solute permeability in the root radius (*K*_*m*_) in various plant hydraulic systems. The model predicted a positive relationship between root pressure and shoot resistance as well as between root pressure and root axial-to-radial hydraulic resistance (*R*_*ra*_*/R*_*rr*_). Based on these predictions, mechanistic studies can be designed to further investigate the role of root pressure in shaping the hydraulic architecture, that is, the whole shoot resistance and the distribution of hydraulic resistances within the plant body. Even if *P*_*r*_ is very low to reach the top of a tall plant, some amount of water will always flow out of the lower leaves when the pressure is large enough to push water to the lower leaf-bearing branches. Our model will be useful in the future to design experiments on root pressure aimed at testing the reliability of predictions presented in this article. Any deviation between the experiment and theory will allow us to modify the theory for improving the predictive performance of the proposed model. The proposed root pressure model is important because it addresses the conditions that may be necessary to reverse the xylem embolism occurring during the day time in plants that generate root pressure at night.

## Acknowledgements

The authors thank Yinshuang Zhang, and Dan Zhou for their assistance with *Bambusa textilis* resistance measurements. This research was supported by grants of the National Natural Science Foundation of China (No. 31770647), the Zhejiang Provincial Natural Science Foundation of China (LY19C150007).

## Supporting information

Additional supporting information may be found in the online version of this article.

Note S1 Derivations for two additions to the theory, namely soil water potential and gravitational potential.

Note S2 The root model output when a simple root is connected to the base of a real bamboo shoot.

Model S1 Excel spreadsheet with root pressure model for bamboo shoot.

